# Aberrant Ca^2+^ homeostasis in adipocytes links inflammation to metabolic dysregulation in obesity

**DOI:** 10.1101/2020.10.28.360008

**Authors:** Ekin Guney, Ana Paula Arruda, Gunes Parlakgul, Erika Cagampan, Nina Min, Yankun Lee, Lily Greene, Eva Tsaousidou, Karen Inouye, Myoung Sook Han, Roger J. Davis, Gökhan S. Hotamisligil

## Abstract

Chronic metabolic inflammation is a key feature of obesity, insulin resistance and diabetes, although the initiation and propagation mechanisms of metaflammation are not fully established, particularly in the adipose tissue. Here we show that in adipocytes, altered regulation of the Ca^2+^ channel inositol triphosphate receptor (IP3Rs) is a key, adipocyte-intrinsic, event involved in the emergence and propagation of inflammatory signaling and the resulting insulin resistance. Inflammation, either induced by cytokine exposure *in vitro* or by obesity *in vivo* lead to increased expression and activity of IP3Rs in adipocytes in a JNK-dependent manner. This results in increased cytosolic Ca^2+^ and impaired insulin action. In mice, adipocyte-specific loss of IP3R1/2 protected against adipose tissue inflammation and insulin resistance despite significant diet-induced weight gain. Thus, this work reveals that IP3R over-activation and the resulting increase in cytosolic Ca^2+^ is a key link between obesity, inflammation and insulin resistance, and suggests that approaches to target adipocyte Ca^2+^ homeostasis may offer new therapeutic opportunities against metabolic diseases, especially since GWAS studies also implicate this locus in human obesity.

## Introduction

The white adipose tissue (WAT) is composed of adipocytes, stromal cells, and immune cells and has potent energy storage, disposal, and endocrine activity, which are critical for the function and survival of the organism(*1–4*). It is well established that nutrient excess and obesity impose chronic pressure and stress on both the storage and endocrine functions of WAT. During chronic overnutrition, adipocytes become increasingly enlarged and dysfunctional and are not able to buffer nutrients effectively, resulting in intracellular stress, recruitment of immune cells, and abnormal production and secretion of inflammatory molecules(*1–5*). Over time, this unfavorable situation ultimately results in tissue dysfunction and damage, systemic metabolic failure including insulin resistance, dyslipidemia, and metabolic disease.

Although the central role of WAT dysfunction in the development of metabolic diseases is well recognized, important questions remain unanswered regarding the mechanisms associated with the emergence and propagation of inflammation in WAT. For example, the specific signals that determine the transition of adipocytes from a healthy to a dysfunctional phenotype during nutritional stress have not been identified, nor it is fully understood how metabolic inflammation is triggered and coupled to metabolic deterioration. At the cellular level, nutrient excess and the expansion of adipocytes leads to organelle stress, particularly in the endoplasmic reticulum (ER). Impaired ER function in the adipocytes is associated with unresolved ER unfolded protein response (UPR), in both mice and humans. However, how ER dysfunction is coupled to WAT inflammation and deterioration is not defined. In fact, in adipocytes, contrary to results in other metabolic cells such as hepatocytes, loss of function of key UPR regulators such as XBP1 (*6*) or ATF6 (*unpublished)* did not affect adipose tissue formation and function under homeostatic metabolic conditions and does not impact systemic metabolism. Thus, ER dysfunction in adipocyte and its impact in cell and systemic metabolism seems to present unique features(*7–11*).

Interestingly, studies performed in *Drosophila melanogaster* show that alterations in mechanisms that control ER Ca^2+^ homeostasis such as loss of function of the IP3R, the ER channel responsible for Ca^2+^ release from ER to the cytosol, leads to obesity and altered lipid metabolism(*12*). Additionally, loss of function of another key mechanism that control ER and cytosolic Ca^2+^ levels, the Store Operated Ca^2+^ Entry (SOCE) in the fat body leads to marked lipid accumulation and obesity in the fly(*13*). Loss of function of SOCE in adipocytes in vitro also leads to alterations in lipid mobilization(*14*). Interestingly, human GWAS studies have shown that an *ITPR* polymorphism is associated with increased body-mass index, fat mass(*15*) and waist-to-hip ratio(*16*) but not metabolic complications and genotype-phenotype causality has yet to be established.

These studies raise the possibility that regulation of intracellular Ca^2+^ homeostasis may be a key mechanism for adipose tissue metabolism. However, whether the regulation of ER Ca^2+^ handling has any impact on adipocyte organelle function or metabolic and inflammatory regulation in mammals have never been explored. Actually, the impact of intracellular Ca^2+^ homeostasis for adipocyte metabolism in general is a largely unexplored area.

In the present study, we identified a critical role for IP3Rs and intracellular Ca^2+^ homeostasis as key drivers of adipocyte stress and adipose tissue inflammation. Through a series of in vitro and in vivo experiments, we demonstrated that IP3R abundance and activity in adipocytes are modulated by inflammatory cytokine exposure in vitro and by high fat diet-induced inflammation in vivo in a mechanism dependent of the kinase JNK. Experimental downregulation of IP3R isoforms in adipocytes decreased inflammatory signaling and metabolic dysfunction in vitro and in vivo, thus demonstrating that regulation of Ca^2+^ homeostasis in adipocytes is a key signal linking metabolic stress to inflammation and metabolic deterioration in obesity.

## Results

### Inflammation activates IP3R and CaMKII phosphorylation through JNK

In order to gain insight into a possible role of intracellular Ca^2+^ regulation in adipocyte dysfunction induced by inflammatory and metabolic stress, we first used an in vitro system where we exposed differentiated 3T3-L1 adipocytes to a low dose of TNF-a, a pro-inflammatory cytokine uniformly increased in both mouse and human obesity(*1*). Addition of TNFα to 3T3-L1 cells led to acute elevation of cytosolic Ca^2+^ (Figure 1A). We then exposed 3T3-L1 cells to TNFα for 6 and 24 hours and assessed the ER Ca^2+^ content in those cells by stimulating ER Ca^2+^ release with the purinergic receptor agonist ATP. Addition of ATP in the absence of extracellular Ca^2+^ lead to decreased cytosolic Ca^2+^ peak in cells previously treated with TNFα for 6h (Figure 1B) and 24h (Supp Figure 1A), suggesting that the TNFα preincubation had already depleted ER Ca^2+^ by shuttling it into the cytoplasm. Consistent with the Ca^2+^ imaging, adipocytes treated with TNFα presented higher CaMKII phosphorylation (Figure 1C). Next, we asked whether changes in adipocyte Ca^2+^ dynamics upon TNFα treatment were related to alterations of ER Ca^2+^ release. ER Ca^2+^ release is coordinated by the action of the IP3Rs and Ryanodine receptors (RyaRs). Upon detecting the predominant presence of IP3Rs compared to RyaRs in adipocytes and adipose tissue (Supp Figure 1B and 1C), we focused on IP3Rs to explore the adipocyte Ca^2+^ homeostasis and its functional impact on metabolism. As shown in Figure 1D and Supp Figure 1D, treatment of adipocytes with TNFα for 6, 12, or 24 hours led to a marked increase in IP3R1 phosphorylation (Ser1756), an indicator of an open channel activity. The increased IP3R1 phosphorylation was also observed after TNFα treatment for longer periods of time up to 4 days (96 hours) (Figure 1F). Thus, TNFα induces elevation in cytosolic Ca^2+^ at least in part by modulating IP3R activity.

**Figure 1:**
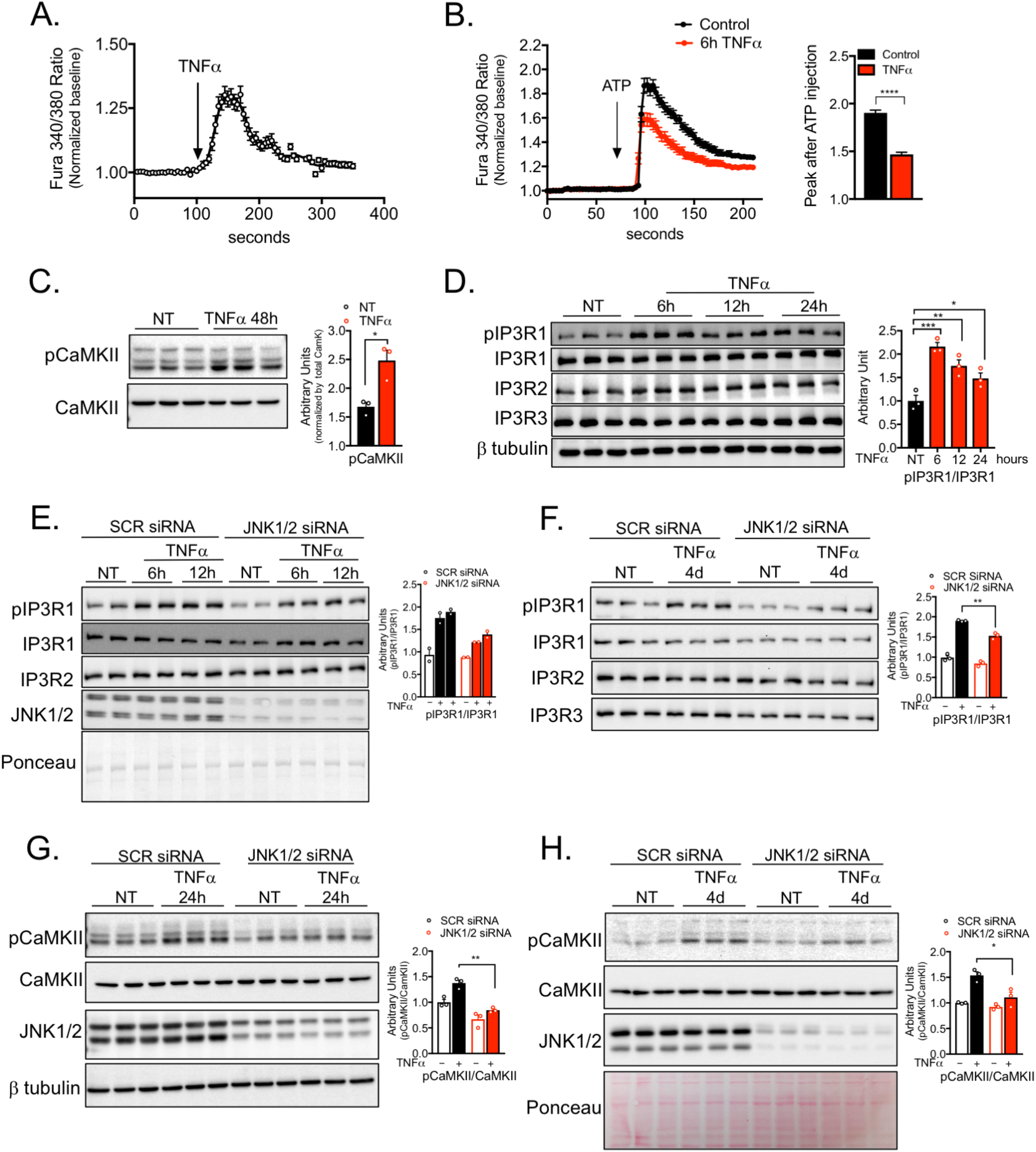
Inflammatory stress leads to increased IP3R expression and activity in adipocytes. (A) Representative Fura-2AM-based cytosolic Ca^2+^ measurements in differentiated 3T3-L1 adipocytes treated with 50ng/mL TNFα. (B) Left panel: representative Fura-2AM-based cytosolic Ca^2+^ measurements in differentiated 3T3-L1 pre-treated with 4ng/mL TNFα for 6 hours. ER Ca^2+^ depletion was stimulated with 50 μM of ATP in media with no extracellular Ca^2+^. Right panel: quantification of Ca^2+^ peak normalized to baseline. n= 405 cells per group, representative of 2 independent experiments, **** p<0.0001. (C) Left panel: Immunoblot analysis of protein expression and phosphorylation levels in 3T3-L1 adipocytes treated with vehicle or 4ng/mL TNFα for 48 hours. Right panel: Quantification of the western blots, n=3, representative of 2 independent experiments, *p=0.016. (D) Left panel: Immunoblot analysis of protein expression and phosphorylation levels in 3T3-L1 adipocytes treated with vehicle or 4ng/mL TNFα for 6,12 and 24 hours. Right panel: Quantification of the western blots n=3, representative of 3 independent experiments, ***p<=0.0004 (6h), **p=0.007 (12h), *p=0.012 (24h). (E) Left panel: Immunoblot analysis of protein expression and phosphorylation levels in 3T3-L1 adipocytes transfected with scrambled (SCR) or JNK1/2 siRNA. Cells were treated with 4ng/mL TNFα or vehicle for 6 and 12 hours. Right panel: Quantification of the western blots, n=2, representative of 2 independent experiments. (F) Left panel: Immunoblot analysis of protein expression and phosphorylation levels in 3T3-L1 adipocytes transfected with scrambled (SCR) and JNK1/2 siRNA. Cells were treated with 4ng/mL TNFα or vehicle for 96h (4d). Right panel: Quantification of the western blots, n=3, ** p=0.0012. (G) Left panel: Immunoblot analysis of protein and phosphorylation levels in 3T3-L1 adipocytes treated with vehicle or 4ng/mL TNFα for 24 hours. Right panel: Quantification of the western blots n=3, **p=0.001. (H) Left panel: Immunoblot analysis of protein and phosphorylation levels in 3T3-L1 adipocytes treated with vehicle or 4ng/mL TNFα for 96 hours (4d). Right panel: Quantification of the western blots, n=3, *p=0.025.

We then investigated the mechanism through which TNFα affects IP3R expression and phosphorylation in adipocytes. Given that JNK is a key inflammatory kinase downstream of inflammatory cytokines involved in metabolic homeostasis(*17–19*), we tested whether JNK mediates the effects of TNFα on IP3R abundance and phosphorylation in adipocytes. For that, we transfected differentiated 3T3-L1 cells with siRNAs against JNK1 and JNK2 or a scrambled siRNA (control) followed by treatment with TNFα for 6, 12 or 96h (4 days). Interestingly, suppression of JNK1/2 lead to reduction in IP3R expression and inhibited TNFα-induced IP3R phosphorylation (Figure 1E and F and Supp Figure 1E). Additionally, we observed that the induction of CaMKII phosphorylation triggered by TNFα in control cells was completely abolished in the absence of JNK1/2 in the 3T3-L1 adipocytes (Figure 1G and 1H). Altogether, these data reveal that activation of inflammatory signaling by TNFα in adipocytes leads to alterations in ER Ca^2+^release and cytosolic Ca^2+^ levels, which leads to activation of Ca^2+^ dependent protein kinase CaMKII, a process downstream of the protein kinase JNK.

### IP3R derived Ca^2+^ release is key for inflammatory and metabolic actions of TNF-a in adipocytes

Next we tested whether IP3R-induced Ca^2+^ release is involved in the inflammatory and metabolic actions of TNFα on adipocytes. For that we performed siRNA-induced knockdown of IP3R1/2/3 in differentiated 3T3-L1 cells. As shown in Figure 2A and Supp Figure 2A, this approach efficiently decreased IP3R1/2/3 expression. Additionally, it significantly inhibited ER-driven Ca^2+^ release stimulated by ATP (Figure 2B). We did not observe any difference in ATP-stimulated cytosolic Ca^2+^ elevation between the downregulation of IP3R1/2 or IP3R1/2/3, thus in the following experiments we continued with combined downregulation of IP3R1/2.

**Figure 2:**
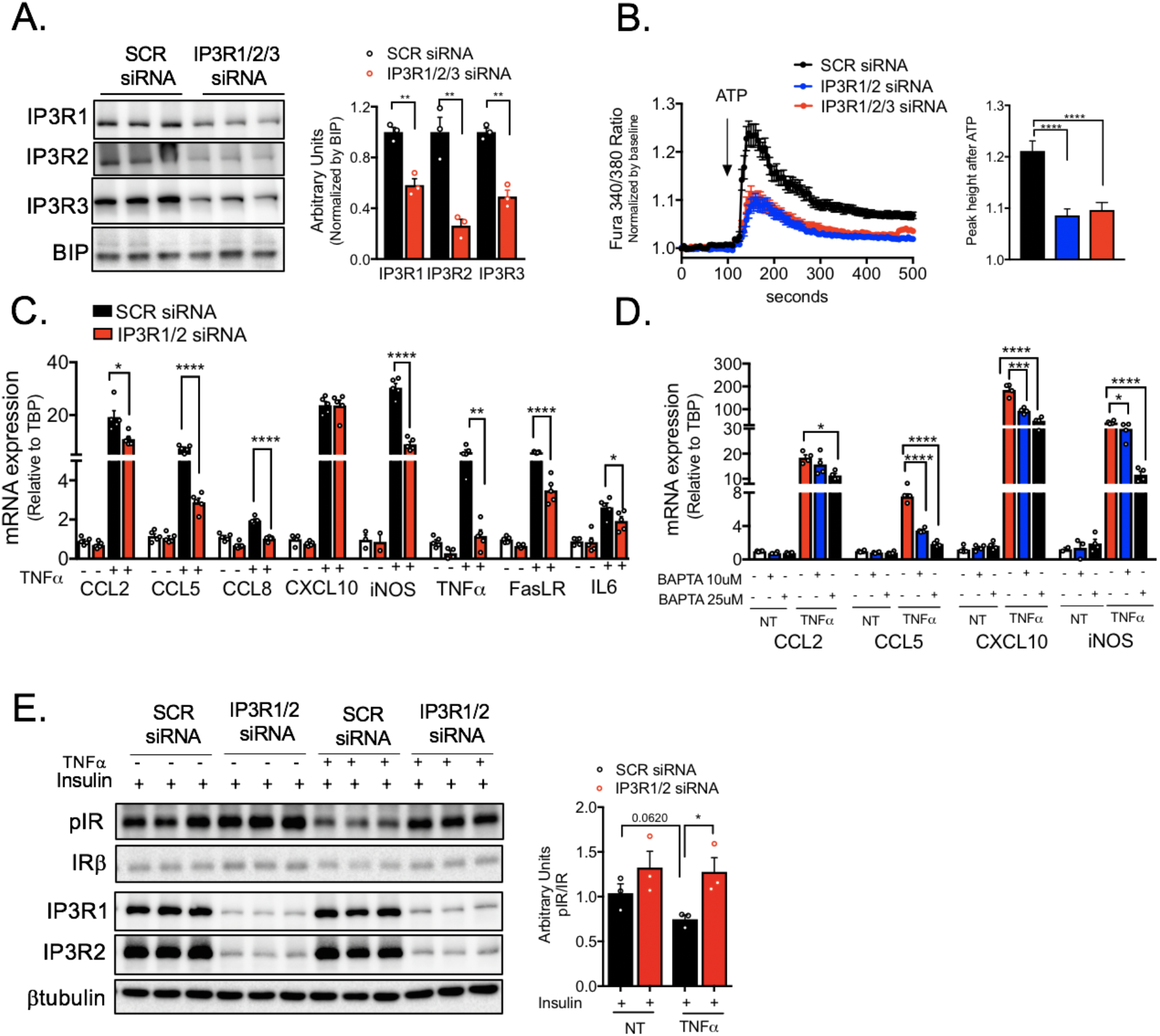
IP3R suppression or chelation of cytosolic Ca^2+^ in adipocytes impairs TNFα inflammatory and metabolic action. (A) Left panel: Immunoblot analysis of protein expression levels in 3T3-L1 adipocytes transfected with scramble (SCR) and IP3R1/2/3 siRNA. Right panel: Quantification of the western blots, n=3, **p=0.029 (IP3R1), **p<0.002 (IP3R2), **p<0.001 (IP3R3). (B) Left panel: Representative Fura-2AM-based cytosolic Ca^2+^ measurements in differentiated 3T3-L1 adipocytes transfected with scramble (SCR), IP3R1/2 of IP3R1/2/3 siRNA. Ca^2+^ release was induced with 50μM of ATP in a media with no extracellular Ca^2+^. Right panel: Quantification of the cytosolic Ca^2+^ peak normalized to baseline, n=236 cells for SCR siRNA and 226 cells for IP3R1/2 and IP3R1/2/3 siRNA. Representative of 2 independent experiments, ****p<0.0001. (C) mRNA levels of the indicated genes evaluated by qPCR derived from 3T3-L1 adipocytes transfected with scramble (SCR) and IP3R1/2 siRNA and treated with 4ng/mL of TNFα or vehicle for 6 hours, n=5, representative of 4 independent experiments. *p<0.014 (CCL2), ****p<0.0001 (CCL5, 8, iNOS, FasL R), ***p=0.0006 (TNF), *p<0.05 (IL6). (D) mRNA levels of the indicated genes evaluated by qPCR derived from 3T3-L1 adipocytes treated with 10 and 25 μM of BAPTA-AM and stimulated with 4ng/mL of TNFα or vehicle for 6 hours, n=4, representative of 2 independent experiments. *p<0.019 (CCL2), ****p<0.0001 (CCL5), ***p=0.0002, ****p<0.0001(CXCL10), *p<0.02,****p<0.0001(iNOS).(E) Immunoblot analysis and quantification of insulin signaling after 3 minutes of 3nM insulin treatment in 3T3-L1 adipocytes transfected with SCR and IP3R1/2 siRNA treated with vehicle or 4ng/mL of TNFα for 96 hours, n=3, representative of 2 independent experiments, * p<0.03.

As shown in Figure 2C, TNFα treatment for 6h led to a marked upregulation in the expression of pro-inflammatory cytokines and chemokines such as CCL2, CCL5, and CCL8, and inflammatory molecules such as iNOS, TNFα itself, and FasL receptor in control adipocytes expressing scrambled (SCR) siRNA. Strikingly, the inflammatory activity of TNFα was markedly diminished in adipocytes where IP3R1/2 were downregulated (Figure 2C). To evaluate whether these effects resulted from changes in ER or cytosolic Ca^2+^ levels upon TNFα exposure, we treated 3T3-L1 cells with BAPTA-AM, a cytosolic Ca^2+^ chelator. BAPTA-AM co-treatment for 6h also significantly reduced TNFα-induced expression of chemokines and inflammatory molecules (Figure 2D and Supp Figure 2B). These data indicate a robust role for IP3Rs in the regulation of TNFα-driven inflammation in adipocytes and provide strong evidence that regulation of cytosolic Ca^2+^ signals have a potent action on inflammatory activity in adipocytes.

It is well known that in adipocytes, chronic low-grade TNFα treatment promotes insulin resistance by a crosstalk between inflammatory and insulin signaling(*5, 20, 21*). We therefore asked whether decreasing IP3R activity by IP3R1/2 knockdown could prevent TNFα-induced insulin resistance in adipocytes. A low-dose TNFα treatment induced marked insulin resistance in control adipocytes, as evaluated by insulin stimulation of insulin receptor (IR) phosphorylation (Figure 2E). Strikingly, the inhibitory effect of TNFα on insulin signaling was abolished in cells lacking IP3Rs 1/2 (Figure 2E). Similar effects were observed in 3T3-L1 cells co-treated with BAPTA-AM and TNFα for 72 hours (Supp Figure 2C) on AKT phosphorylation, indicating that elevation in cytosolic Ca^2+^ underlies both the inflammatory and metabolic activity of TNFα.

### Obesity leads to increased IP3R content and activity in adipose tissue

We next explored the potential role of IP3Rs in adipose tissue dysfunction in vivo and in obesity. For that, we first examined the impact of obesity on the expression levels of all three IP3R isoforms in epididymal white adipose tissue lysates (eWAT). As shown in Figure 3A and Supp Figure 3A, the protein and mRNA expression levels of IP3R1-3 were markedly upregulated in mice fed a high-fat diet (HFD). Along with higher expression, obesity led to enhanced IP3R1 phosphorylation (Ser1756) (Figure 3A). Because the eWAT derived from obese animals contains large number of immune cells, we asked whether the presence of IP3R channels reflected expression in infiltrating cells or is intrinsic to adipocytes. To address this possibility, we fractionated the adipose tissue from lean (low-fat diet) and HFD fed mice in order to separate the stromal vascular fraction (SVF) from the buoyant adipocyte layer. Examination of adiponectin as a specific adipocyte marker confirmed the validity of the fractionation (Supp Figure 3B). As shown in Figure 3B and C, the increased expression of IP3R1/2 in HFD-fed mice was derived from the adipocytes, with no significant differences detected in the SVF fraction. We additionally evaluated the expression of IP3Rs in epididymal adipose tissue derived from a genetic model of obesity, the leptin-deficient mice (*ob/ob*). Similar to the profile observed in HFD, eWAT derived from ob/ob mice also presented strikingly higher expression of IP3R1 and IP3R2 but no significant differences in IP3R3 content (Figure 3D). To determine whether the increased quantity and activity of IP3Rs in adipocytes affects cytosolic Ca^2+^ levels, we measured the phosphorylation status of CaMKII as a proxy for cytosolic Ca^2+^ (*22, 23*). We were not able to obtain consistent reads when trying to measure cytosolic Ca^2+^ levels in adipose tissue in vivo with the available Ca^2+^ imaging reporters. As shown in Figure 3E, in mice with both dietary (HFD, upper panel) and genetic (lower panel) obesity, phosphorylation of CaMKII in eWAT was significantly elevated compared with their lean counterparts. These data strongly support that obesity leads to altered Ca^2+^ homeostasis in adipocytes, at least in part, as a result of higher activity of the IP3Rs.

**Figure 3:**
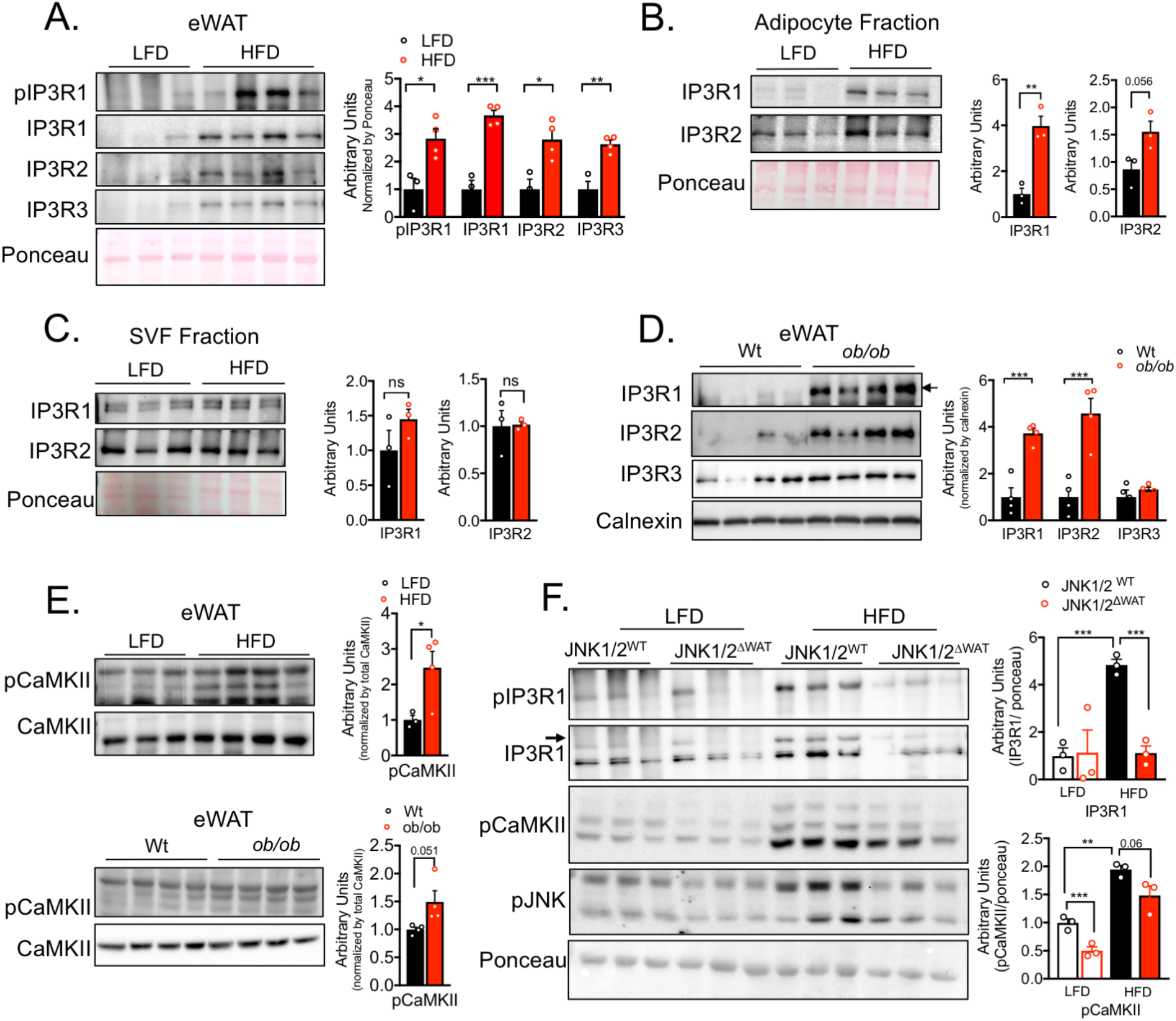
Obesity leads to increased IP3R expression and pCaMKII phosphorylation in adipose tissue. (A) Left panel: Immunoblot analysis of protein expression and phosphorylation levels in epididymal white adipose tissue (eWAT) from low fat diet (LFD) and high fat diet (HFD)-fed mice (16 weeks). Right panel: Quantification of the western blots, n=3 for LFD and 4 for HFD, representative of 3 independent experiments. *p<0.017, **p< 0.0027, ***p< 0.0006. (B and C) Left panel: Immunoblot analysis of protein expression in adipocyte fraction and stromal-vascular fraction (SVF) derived from epididymal white adipose tissue (eWAT) from mice fed low fat diet (LFD), n= 3 or high fat diet (HFD) for 16 weeks, n=3. Right Panel: Quantification of the western blots n=3 per group, representative of 2 independent experiments, **p<0.001. (D) Left panel: Immunoblot analysis of protein expression and phosphorylation levels in epididymal white adipose tissue (eWAT) of Wt and leptin-deficient (*ob/ob)* mice. Right panel: Quantification of the western blots n=4 per group, representative of 2 independent experiments, *p=0.0009. (E) Immunoblot analysis of protein expression and phosphorylation levels of CaMKII in epididymal white adipose tissue (eWAT). Upper panel, low fat diet (LFD) n= 3 and high fat diet (HFD) for 16 weeks n=4. Lower panel: Wt and leptin-deficient (*ob/ob)* mice n=4 per group. Right panel: Quantification of the western blots, *p<0.045. (F) Left panel: Immunoblot analysis of protein expression and phosphorylation levels in eWAT derived from controls (JNK1/2^WT^) and adipocyte-specific loss of JNK1/2 (JNK1/2^ΔWAT^) fed an LFD or HFD for 16 weeks. Right panel: Quantification of the western blots n=3 per group, *** p<0.0005 (IP3R1), **p=0.007, ***p=0.0007 (pCaMKII). In all panels error bars denote s.e.m.

In our experiments with cultured adipocytes, we found that enhanced IP3R expression and activity upon TNFα treatment was dependent on the presence of JNK1/2. It is critical to examine this paradigm in vivo, in the adipose tissue. Hence, we measured IP3R expression and phosphorylation levels in eWAT derived from mice with adipocyte-specific loss of both JNK1/2. As shown in Figure 3F and Supp Figure 3C and 3D, total and phosphorylated IP3R1 are significantly decreased in eWAT from JNK1/2 deficient mice maintained on HFD for 16 weeks. IP3R1 and IP3R2 downregulation in mice with adipocyte-specific JNK1/2-deficiency was also observed at mRNA levels (Supp Figure 3E). Interestingly, we also determined that the increased phosphorylation of CaMKII promoted by HFD was diminished in the eWAT from mice deficient in JNK1/2 in adipocytes (Figure 3F). These results strongly support a novel and previously unrecognized role for JNK signaling in modulating IP3R abundance and activity in vitro and in vivo and also suggests that the increased IP3Rs expression levels detected in obese adipose tissue is likely a consequence of an inflammatory activation of JNK and serves to integrate critical mechanistic nodes linking obesity and adipose tissue inflammation in vivo.

### Loss of IP3R1/2 in adipose tissue protects from high fat diet induced inflammation

To investigate the impact of IP3Rs in adipocyte tissue metabolism in vivo, we generated mice with adipocyte-specific loss of function of IP3R1 and 2 (IP3R1/2^AdpCre^) by crossing mice with IP3R1 and IP3R2 (IP3R1/2) floxed alleles with mice expressing Cre recombinase under the control of Adiponectin promoter (Adp-Cre) (Supp Figure 4A). Expression of Cre recombinase in adipocytes from IP3R1/2^fl/fl^ significantly decreased IP3R1 and 2 protein and mRNA levels as shown in multiple depots, with no significant differences in the expression level of IP3R3 (Figure 4A and Supp Figure 4B and C). We then evaluated the impact of adipose tissue specific IP3R1/2 loss of function on body weight gain and energy expenditure in mice fed a low-fat diet (LFD) or 60% high fat-diet (HFD). As shown in Supp Figure 4D, on a LFD, body weight gain between genotypes was indistinguishable. However, after 12 weeks on HFD the body weight gain between the groups diverged, and by 16 weeks, IP3R1/2^AdpCre^ mice were 15-20% heavier than control mice (Figure 4B). DEXA analysis determined that the difference in body weight at 16 weeks of HFD was largely due to higher fat mass accumulation in IP3R1/2^AdpCre^ mice, and no differences were observed in lean mass between the genotypes (Figure 4C). IP3R1/2^AdpCre^ mice also showed a tendency for decreased VO2 and VCO2 (Supp Figure 4E and F), especially after the addition of the β3-agonist CL-316,243. No differences in body weight or food intake were observed at this time point (Supp Figure 4G and H). Although intracellular Ca^2+^ regulation can influence lipolysis, we did not see any significant differences in basal and isoproterenol (Iso) stimulated lipolysis in these mice (Supplemental Figure 4I and J). These data indicate that IP3R1/2-deficiency leads to susceptibility to obesity and increased fat mass, similar to what has been observed in fruit-flies(*12*) and in GWAS studies(*15, 16*).

**Figure 4:**
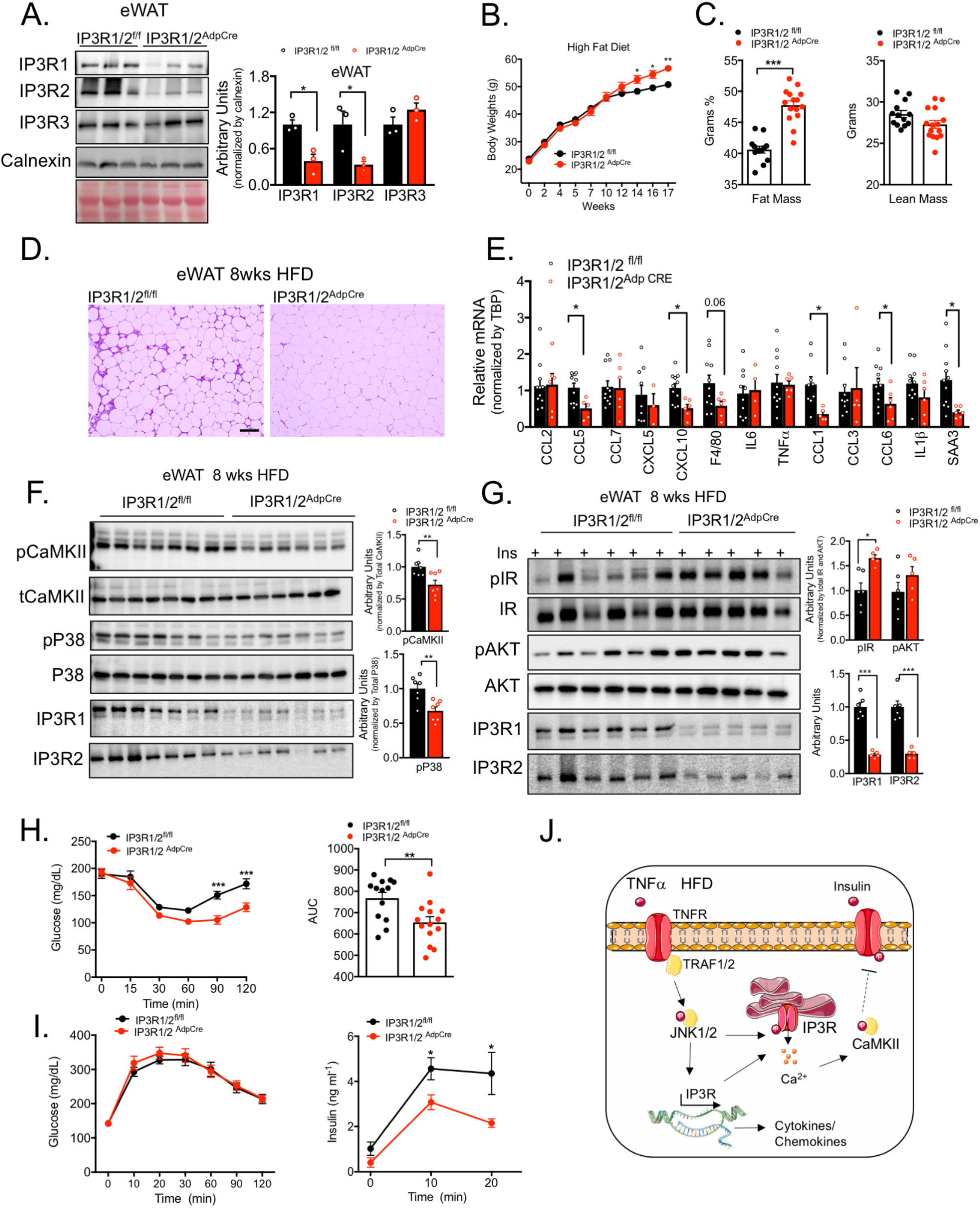
IP3R1/2 deficiency leads to increased weight gain and decrease adipose tissue inflammation. (A) Left panel: Immunoblot analysis of protein expression levels epididymal white adipose tissue (eWAT) from control (IP3R1/2^fl/f^ mice) and adipocytespecific loss of function of IP3R1/2 (IP3R1/2^AdpCRE^). Animals were fed chow diet. Right panel: Quantification of the western blots, n=3 per group, eWAT: * p<0.01 (IP3R1), *p<0.04 (IP3R2) (B) Weight gain curves of control (IP3R1/2^fl/f^ mice) and adipocyte-specific loss of function of IP3R1/2 (IP3R1/2^Ad^P^CRE^) mice on HFD, n=13 for IP3R1/2^fl/f^ and n=15 for IP3R1/2^AdpCRE^.Representative of 3 independent cohorts, *p<0.05,** p=0.001. (C) DEXA analysis of whole-body fat and lean mass, n= 13 for IP3R1/2^fl/f^ mice and n=15 for IP3R1/2^AdpCRE^, ***p<0.0001. (D) Representative hematoxylin and eosin stained histology section derived from epidydimal white adipose tissue (eWAT) from weight matched control (IP3R1/2^fl/f^ mice) and adipocyte-specific loss of IP3R1/2 (IP3R1/2^AdpCRE^) fed on HFD for 8-9 weeks. (E) eWAT mRNA levels of the indicated genes evaluated by qPCR of control (IP3R1/2^fl/f^ mice) and adipocyte-specific loss of function for IP3R1/2 (IP3R1/2^AdpCRE^) mice fed HFD for 8-9 weeks, n= 11 for IP3R1/2^fl/f^ and n=6 for IP3R1/2^AdpCRE^ *p<0.012 (CCL5), *p<0.007 (CXCL10), *p<0.016 (CCL1), *p<0.04 (CCL6), *p<0.015 SAA3. (F) Left panels: Immunoblot analysis of protein expression and phosphorylation levels in epididymal white adipose tissue (eWAT) lysates from control (IP3R1/2^fl/f^ mice) and adipocyte-specific loss of function of IP3R1/2 (IP3R1/2^AdpCRE^) fed on HFD for 8-9 weeks. Right panel: Quantification of the western blots, n=7 per group, ** p<0.04. (G) Left panel: Markers of insulin signaling evaluated by immunoblot analysis of total eWAT from animals injected with insulin (0.45 U/kg) through the inferior vena cava. Tissues were collected 3 min after injection. Right panel: Quantification of phosphorylation levels of indicated proteins normalized to total protein levels, n= 6 for IP3R1/2^fl/f^ and n=5 for IP3R1/2^Ad^p^CRE^ * p<0.004, *** p<0.0001. (H) Left panel: Insulin tolerance test in control (IP3R1/2^fl/f^ mice) and adipocyte-specific loss of function of IP3R1/2 (IP3R1/2^Ad^p^CRE^) mice fed HFD for 16 weeks, ***p=0.0004. Right panel: Quantification of area under the curve, n= 13 for IP3R1/2^fl/f^ and n= 14 for IP3R1/2^AdpCRE^, representative of 2 independent experiments, ** p<0.006. (I) Left panel: Oral glucose tolerance test in control (IP3R1/2^fl/fl^ mice) and adipocyte-specific loss of function of IP3R1/2 (IP3R1/2^Ad^p^CRE^) mice fed HFD for 10 weeks, n= 13 for IP3R1/2^fl/f^ and n= 15 for IP3R1/2^AdpCRE^ animals, representative of 2 independent experiments. Right panel: Insulin levels during the oral glucose tolerance test. *p<0.05 (10min), **p=0.0032 (20min). (J) Working model. In all panels error bars denote s.e.m.

We next examined the effect of IP3R1/2 downregulation on adipose tissue inflammation and stress. In mice fed with a LFD, there were no differences between genotypes in the inflammatory status of WAT (Supp Figure 5A). However, obese IP3R1/2^AdpCre^ mice were markedly protected from eWAT immune infiltration induced by 8-9 weeks of HFD feeding (Figure 4D and Supp Figure 5B), a time when the body weights of both genotypes were similar. In addition, the expression of inflammatory molecules CCL1,5,6, CXCL10, F4/80 and SAA3 were significantly reduced in eWAT tissues derived from IP3R1/2^AdpCre^ compared to controls (Figure 4E and Supp Figure 5C). Liver steatosis was also significantly decreased in IP3R1/2^AdpCre^ mice compared with controls (Supp Figure 5D). We then investigated the impact of IP3R1/2 loss on high-fat diet induced phosphorylation of CaMKII and of stress kinases downstream of inflammation such as JNK, p38, pERK1/2. As shown in Figure 4F, IP3R1/2 loss of function in adipose tissue lead to significantly decreased levels of pCaMKII, suggesting that deletion of IP3R1/2 channels resulted in decreased adipocyte cytosolic Ca^2+^ levels in vivo. Additionally, we detected that loss of IP3R1/2 also significantly inhibited phosphorylation of p38 (Figure 4F) and ERK (supp Figure 5E) with no significant effect difference on phosphorylation of JNK (supp Figure 5E). Notably, this protection from inflammatory stress preceded the development of body weight differences between the groups (Figure 4B). At the end of the 16 weeks of HFD, expression of CCL2 and CCL5 remained significantly lower in the eWAT of IP3R1/2^AdpCre^ mice than controls while other inflammatory changes subsided (Supp Figure 5F).

### Loss of function of IP3R1/2 in adipose tissue protects against insulin resistance

Adipose tissue inflammation is closely associated with insulin resistance. Given the impact on IP3R1/2 deletion on decreasing inflammation and stress signaling in adipose tissue, we investigated the effect of IP3R1/2 deficiency in tissue and systemic glucose metabolism. On a low-fat diet, as with all other parameters measured, IP3R1/2 deletion in adipose tissue had no effect on glucose tolerance and insulin sensitivity (Supp Figure 6A and B). After five weeks of HFD, IP3R1/2^AdpCre^ mice exhibited improved insulin sensitivity at the tissue level (Supp Figure 6C), but this was not sufficient to alter glucose or insulin tolerance (Supp Figure 6D and E). At later times on HFD (9 weeks), IP3R1/2^AdpCre^ mice exhibited improved insulin signaling in eWAT compared to IP3R1/2^fl/fl^ mice, as demonstrated by a significant increase in pIR and a trend towards increased pAKT (Figure 4G). Accordingly, the IP3R1/2^AdpCre^ mice were more insulin sensitive when challenged with an insulin tolerance test (Figure 4H). During an oral glucose tolerance test, the glucose excursion curves did not differ between IP3R1/2^fl/fl^ and IP3R1/2^AdpCre^. However, the glucose-induced insulin secretion in IP3R1/2^AdpCre^ mice was significantly lower than in control mice (Figure 4I), indicating increased insulin sensitivity. Thus overall, adipose tissue-specific loss of function of IP3R1/2 increased diet-induced weight gain but protected against adipose tissue inflammation and insulin resistance and fatty liver development, thereby representing a metabolically healthy obese phenotype.

## Discussion

The discovery of the immunometabolic nature of obesity and the central role of metabolic inflammation in adipose tissue dysfunction in obesity and diabetes have been followed up by a large amount of research connecting the immune response to metabolism and metabolic disease(*1, 5, 20, 24, 25*). However, the exact molecular events linking obesity-driven intracellular stress to the development and propagation of inflammation and subsequent metabolic pathologies have remained elusive and complex. Here, we show that dysregulation of intracellular Ca^2+^ homeostasis is a key, previously unrecognized, link integrating these mechanistic processes into a cohesive model of pathogenesis. Specifically, we show that inflammation, modelled in cells in vitro or during the course of obesity in vivo, leads to enhanced expression and activation of the ER Ca^2+^ channel IP3R in adipocytes, leading to leakage of ER Ca^2+^ into the cytoplasm and activation of the CaMKII, which has important implications in inflammatory signaling and the development of insulin resistance.

Interestingly, the cycle of increased IP3R1 expression and activity promoted by inflammatory stress and chronic overnutrition is prevented in the absence of JNK1/2, suggesting that JNK is a key upstream factor regulating IP3R induced Ca^2+^ release and CaMKII phosphorylation in adipocytes (Figure 4J), which has not been possible to show in a metabolic target and in vivo context. While a previous report suggests that JNK regulates IP3R1 expression at a transcription level in a neuronal cell line(*26*), it is possible that JNK also regulates IP3R1 activity through other mechanisms such as post-translational modification. It is well established that mice harboring JNK deficiency in adipose tissue are protected against metabolic disease, even when challenged by a high fat diet(*27*). Our data suggest that this protection may, at least in part, be mediated by decreased IP3R-driven Ca^2+^ release in JNK1/2 deficient adipocytes.

Inhibition of the rise in cytosolic Ca^2+^, either by downregulation of IP3R expression or treatment with a Ca^2+^ chelator, BAPTA, significantly diminishes the inflammatory cascades triggered by TNFα in cultured adipocytes and adipose tissue inflammation and immune cell infiltration promoted by HFD in vivo. An important impact of the attenuation of inflammatory signaling by downregulation of IP3R1/2 or BAPTA treatment was a striking rescue of the insulin resistance caused by inflammatory stress in vitro and nutritional stress in vivo, implicating regulation of insulin action by cytosolic Ca^2+^. Inhibition of TRPV4, a non-selective Ca^2+^ channel, has also been reported to reduce adipose tissue inflammation(*2δ*) and cytosolic Ca^2+^ has been linked to insulin signaling through an CaMKII-P38-ATF6 axis in liver(*29*). In our model, CaMKII and p38 phosphorylation were both decreased by IP3R1/2 downregulation in adipose tissue, suggesting that the crosstalk between cytosolic Ca^2+^ and insulin signaling in adipose tissue may be mediated by these kinases (Figure 4J). It is possible that other mechanisms regulated by Ca^2+^ contribute to this phenomenon, such as alterations in mitochondrial Ca^2+^ and oxidative stress. Additional work will be necessary to understand how IP3R modulation affects mitochondrial Ca^2+^ levels and function in adipose tissue.

In summary, our work demonstrates that proper regulation of intracellular Ca^2+^ homeostasis is a central process for the maintenance of adipocyte metabolic health, and alterations of this fine balance is a signal that leads to adipocyte dysfunction in the context of obesity. Together with the fact that dysregulated IP3R content and activity in the liver is involved in altered hepatic glucose production(*30–33*), the findings demonstrated here suggest that imbalanced intracellular Ca^2+^ at the IP3R level is a major pillar of metabolic homeostasis and alterations in its function collectively predispose to overall metabolic defects(*29–33*). Moreover, this mechanism seems to be well conserved across different organisms, since in *Drosophila* loss of function of IP3R(*12*) and SOCE(*13*) in the fat body leads to increased adiposity. IP3R1/2 deficiency in the adipose tissue, leads to development of greater adiposity uncoupled of metabolic complications, possibly representing a model of metabolically healthy obesity. Therefore, our work supports the idea that preventing aberrant IP3R activity could be a powerful strategy to rescue systemic glucose metabolism and inflammatory stress in obesity. Indeed, pharmacological attempts in this direction are being developed such as the use of SERCA agonists(*34*) or azoramide(*35*), although it remains to be elucidated whether targeting this mechanism can safely improve metabolism in humans.

## Methods

### General animal care and study design

All *in vivo* studies were approved by the Harvard Medical Area Standing Committee on Animals. Unless stated otherwise, mice were maintained from 4-20 weeks of age on a 12-hour-light /12-hour-dark cycle in the Harvard T.H. Chan School of Public Health pathogen-free barrier facility with free access to water and to a standard laboratory chow diet (PicoLab Mouse Diet 20 #5058, LabDiet). The sample size and number of replicates for this study were chosen based on previous experiments performed in our lab and others(*31, 36*).

### Animal models of obesity

Leptin deficient *Lep^ob^ (ob/ob)* mouse: Wild type, heterozygous and *Lep^ob^ (ob/ob)* mice (Stock no. 000632) were purchased from Jackson Laboratories at 6-7 weeks of age and used for experimentation between 8-10 weeks of age. Diet-induced obesity: Male C57BL/6J mice were purchased from Jackson Laboratories and placed on HFD (D12492: 60% kcal% fat; Research Diets) for up to 20 weeks. Control mice of the same age were fed with a low-fat diet (PicoLab Mouse Diet 20 #5053, LabDiet).

### IP3R1/2 adipocyte-specific deficient mice

Mice carrying floxed alleles for IP3R1 and IP3R2 (Figure 4A) were kindly provided by Dr. Andrew Marks from Columbia University Medical Center(*37, 38*). To generate adipocyte-specific IP3R1/2 deficient mice, IP3R1 and IP3R2^fl/fl^ mice were bred to C57BL/6J mice expressing CRE recombinase under the control of the adiponectin promoter (Adp-Cre, Jax stock #028020). Adp-Cre-mediated recombination of floxed IP3R1/2 alleles was detected in genomic DNA by PCR performed with the following protocol: 3 minutes at 95°C, 35 times repeats of (30 seconds at 95°C, 30 seconds at 60°C, 45 seconds at 72°C), 3 minutes at 72°C and hold at 4°C. Products of PCR were as follows; IP3R1-WT: 320bp; IP3R1-floxed: 406bp; IP3R2-WT: 248 bp, IP3R2-floxed: 394 bp. The genetic background of the mice was determined to be 93-98% C57BL/6. Age-matched littermates were used for the study. For the high fat diet (HFD) studies, mice were placed on HFD (D12492: 60% kcal% fat; Research Diets) from 6 up to 20 weeks. The control chow group were switch from (PicoLab Mouse Diet 20 #5058, LabDiet) to low-fat chow diet (PicoLab Mouse Diet 20 #5053, LabDiet) at 6 weeks of age and remained on this diet the same period than the HFD.

### JNK1/2 adipocyte-specific deficient mice

Mice carrying floxed alleles for JNK1 and JNK2 were generated in Dr. Roger Davis’s laboratory at the Program in Molecular Medicine, UMASS Medical School. JNK1/2 wild type mice crossed with mice carrying Cre recombinase under the control of Adiponectin promoter was used as controls. JNK1/2 flox/flox mice crossed with mice carrying Cre recombinase under control of Adiponectin promoter was used to delete JNK1/2 specifically in adipocytes. Age-matched mice were used for the study. For the high fat diet (HFD) studies, mice were placed on HFD for up to 16 weeks. LFD was Purina Cat# IsoPro3000 and HFD was Bioserv # S3282.

### Mice respiration

Mouse oxygen consumption and carbon dioxide emission were measured using a Columbus Instruments Oxymax-Comprehensive Lab Animal Monitoring System (CLAMS) system, according to guidelines for measuring energy metabolism in mice(*39*). We measured brown adipose tissue-mediated respiration by intrapertoneal injection of the β3 adrenergic receptor-specific agonist, CL316,243 (CL, Tocris, 0.5 mg/kg in 0.9 % w/v in NaCl).

### Glucose and insulin tolerance tests

Glucose tolerance tests: Mice were administered glucose by oral gavage after overnight fasting (lean: 1.5 g kg^-1^, obese: 0.5-1.0 g kg^-1^), and blood glucose levels were measured throughout 120 minutes as indicated in the figures. For insulin tolerance tests, insulin was injected intraperitoneally (0.75-1U/ kg^-1^) after 6h food withdrawal and blood glucose levels were measured throughout 120 minutes as indicated in the figures.

### In vivo lipolysis

Mice were injected intraperitoneally with isoproterenol (10mg/kg in PBS, Tocris) after 6h food withdrawal. Blood was collected by tail vein throughout 60 minutes as indicated in the figure. Free fatty acids and glycerol were determined in the plasma.

### Cell culture

For all in vitro experiments 3T3-L1 cells (ATCC) were seeded into culture dishes at approximately 70% confluence in DMEM with 4.5mM glucose (GIBCO) supplemented with 10% bovine calf serum (CCS) and 1% penicillin/streptomycin. For differentiation, once the cells reached 100% confluence (2-3 days after seeding), medium was changed to DMEM with 10% fetal bovine serum (FBS) with 1% penicillin/streptomycin. After 2 days (day 0), adipocyte differentiation was induced by the addition of 500 μM 3-isobutyl-1-methylxanthine (IBMX), 5 μg/ml insulin, 10 μM dexamethasone, and 10 μM rosiglitazone. On day 2 medium was switched to DMEM with 10% FBS and 1% penicillin/streptomycin, 5 μg/ml insulin and 10 μM rosiglitazone. On day 4, medium was switched to DMEM with 10% FBS and 1% penicillin/streptomycin (maintenance medium), and this medium was replaced every other day until day 10, when the adipocytes were fully differentiated. The experiments were performed at day 10.

### TNFα and BAPTA-AM treatment

Recombinant murine-TNFα (cat number: 315-01A, PeproTech) was dissolved in nuclease-free sterile water and diluted according to the user’s manual. The main stock was prepared at concentration of 100 ug/mL and aliquoted to prevent thawing-refreezing. For all experiments, TNFα was diluted to 4 ng/ml in the maintenance medium and added to cells for the period of time described in the experiments at 37°C. After TNFα treatment, the medium was removed, wells were washed with PBS and the tissue culture plates were frozen in liquid nitrogen. For experiments involving BAPTA-AM (cat number: A1076, Sigma-Aldrich), main stocks were prepared at 10 mM in DMSO according to user’s manual and diluted to 1 uM or 10 uM in maintenance medium. The amount of DMSO in the BAPTA-AM solution was added to TNFα-only and control medium treatment mixtures. For TNFα treatment experiments, treatment mixtures were prepared as TNFα-only (with DMSO), TNFα with BAPTA-AM, BAPTA-AM-only, and control medium (with DMSO). Treatment mixtures were added simultaneously, and culture plates were incubated at 37°C.

### Transfection protocol

For the IP3R1/2/3 and JNK1/2 knockdown experiments, 10-day-diffentiated adipocytes were transfected with scramble or IP3R1/2/3 siRNA (separately or in combination) or JNK1/2 siRNA. For the transfection, cells were trypsinized (0.25% trypsin), transferred into 15 ml conical tubes and centrifuged for 10 minutes at 300 rpm at 4°C. The supernatant was removed, and the pellets were diluted in transfection media (DMEM, 10%FBS, no penicillin/streptomycin). The adipocytes were counted and seeded in 6 or 12-well plates (5×10^5^ cells/well for 6-well plates and 2.5×10^5^ cells/well for 12-well plates; at approximately 70% confluence). Lipofectamine-RNAiMax (Catalog number: 13778030) and Opti-MEM medium were from Thermo Fisher Scientific; Mouse-siRNAs for On-target SMARTpool scrambled-control, ITPR1 (catalog ID: L-040933-00-0005), ITPR2 (L-041018-00-0005), ITPR3 (L-065715-01-0005), Mapk8 (L-040128-00-0005) and Mapk9 (L-040134-00) were from Dharmacon. Transfection mixtures were prepared following the lipofectamine-RNAiMAX-transfection protocol. The transfection mixture (300ul/well for 6-well plate) was added to the wells containing adipocytes, bringing the total volume of the wells up to 2.2 ml. The culture plates were incubated in 37°C incubators for 16 hours. After 16 hours, medium was replaced with maintenance medium and plates were incubated until 36 hours after the transfection before performing the experiments.

### Insulin signaling experiments

3T3-L1 cells transfected with scrambled or IP3R1/2 siRNA were treated with 4ng/mL of TNFα for 4 days with or without BAPTA-AM. Media was changed every 24-hours. After 96 hours (4 replacements), cells were washed with warm (37°C) PBS three times. Then, serum-free medium was added into the wells (6-well plates) for 6 hours at 37°C. Insulin solutions were prepared in serum-free medium (1, 10 and 100 nM). After 6 hours, the cells were treated with serum-free medium with or without 3nM insulin solution for 3 minutes. After insulin treatment wells were washed twice with cold PBS and frozen in liquid nitrogen.

### Total protein extraction and western blotting

Adipose tissues were homogenized in cold lysis buffer containing 50 mM Tris-HCl (pH 7.4), 2 mM EGTA, 5 mM EDTA, 30 mM NaF, 10 mM Na_3_VO_4_, 10 mM Na_4_P2O_7_, 40 mM glycerophosphate, 1% NP-40, and 1% protease inhibitor cocktail or RIPA buffer (Cell signaling #9806) supplemented by 1% protease inhibitor cocktail. Adipose tissue was homogenized using a polytron in cold lysis buffer. Homogenates were placed on ice for 45 minutes and during this time intermittent vortexing was performed. The homogenates were centrifuged for 15 minutes at 9000 rpm to pellet cell debris. The supernatant with the protein lysate was collected and the centrifugation was repeated once more. Protein concentrations were determined by BCA. Finally, samples were diluted in 5x Laemmli buffer and boiled for 5 minutes 95°C. The protein lysates were subjected to SDS-polyacrylamide gel electrophoresis, as previously described(*31, 35*). Membranes were incubated with anti-JNK (Cell signaling #9252), anti-pAKT-Rabbit Antibody(Ser473) (Cell Signaling #9271), anti-AKT-Rabbit Antibody (Cell Signaling #9272),anti-IR (Santa Cruz 711), p-Insulin Receptor Antibody(p-Thy1162/1163) (Calbiochem 407707 or Sigma I1783), phospho-CaM KinaseII (Cell Signaling #12716), CamKII (pan) (Cell Signaling #3362), pIP3R1 (Cell signaling #3760), IP3R1 (Bethyl Laboratories – A392-158A), IP3R2 (received as a kind gift from Dr Wojcikiewicz, Upstate Medical University), purified mouse anti-IP3R3 (BD Transduction Laboratories), BiP (Cell Signaling #3183), Calnexin (Santa Cruz, sc-6465), Adiponectin-Acrp30 (G-17 clone,sc-26496), β-tubulin (Abcam, ab 21058).

### SVF and adipocyte isolation

Epididymal adipose tissue was collected in PBS, washed twice with fresh PBS and transferred into 10 cm petri-dishes on ice containing enzymatic digestion buffer (Krebs-Ringer solution, 2% BSA, Collagenase type-I (1.25mg powder/1ml); 4-5 ml/mouse for lean mice and 7-8 ml/ HFD mice). Immediately after, tissue was minced using surgical scissors for 5-10 minutes until each piece is smaller than 1 mm. The mixture of minced adipose tissue and enzymatic digestion buffer was transferred into 50 ml conical tube and placed on ice. Once this procedure was completed for each mouse, the mixtures were transferred into a shaker and incubated for 45-60 minutes at 37°C at 100 rpm until all adipose tissue fragments were digested completely. Digested mixtures were filtered using 200um strainers (pluriStrainer 200 μm (Cell Strainer); Catalog ID: 43-50200-03). For adipocyte fraction preparation, the supernatant adipocyte fraction was collected and transferred into 15 ml conical tubes which was then brought up to 10 mL volume by the addition of Krebs-ringer-2%BSA-mixture followed by centrifugation at 300 rpm for 10 min at 4°C. Following centrifugation, supernatant adipose tissue fraction was transferred into 1.5 ml tubes, which were then centrifuged at 300 rpm for 10 min. Infranatant buffer was then removed, lysis buffer was added, and the protein lysate was prepared as described for whole tissue. For stromal-vascular fraction (SVF) preparation, the buffer between the supernatant and the pellet was removed and the remaining pellet was mixed with PBS and centrifuged at 3000 rpm for 5 minutes at 4°C. Following centrifugation, the supernatant buffer was removed, lysis buffer was added to the remaining pellet and protein lysate was prepared as described for whole tissue.

### Gene expression analysis

Tissues were homogenized in Trizol (Invitrogen) using TissueLyser (Qiagen). To obtain RNA, Trizol homogenates were mixed with chloroform, vortexed thoroughly and centrifuged at 12000g for 20 min at 4°C. The top layer was transferred to another tube and mixed with isopropanol and centrifuged again at 12000g for 20 min at 4°C. The RNA in the precipitate was washed twice with 70% ethanol and diluted in RNAse free water. Complementary DNA was synthesized using iScript RT Supermix kit (Biorad). Quantitative real-time PCR reactions were performed in duplicates or triplicates on a ViiA7 system (Applied Biosystems) using SYBR green and custom primer sets or primer sets based on Harvard primer bank. Gene of interest cycle thresholds (Cts) were normalized to TBP housekeeping levels by the ΔΔCt. For Supplemental figure 3E, mRNA expression was examined by quantitative RT-PCR analysis using a Quantstudio machine (ThermoFisher Scientific). TaqMan assays were used to quantify IP3R1 (Itpr1, Mm00439907_m1), IP3R2 (Itpr2, Mm00444937_m1), and IP3PR3 (Itpr3, Mm01306070_m1) mRNA (ThermoFisher Scientific).

### Cytosolic Ca^2+^ imaging using Fura-2 AM

Cells were loaded with 4 μM Fura-2AM and 1 μM Pluronic F-127 in HBSS for ~30 to 60 min at room temperature. Before imaging, the cells were washed and kept in a medium containing 10 mM HEPES, 150 mM NaCl, 4 mM KCl, 2 mM CaCl_2_, 1 mM MgCl_2_, 10 mM D-glucose, pH 7.4 for 10 minutes. Ca^2+^-free medium was prepared similarly to the buffer described above, in the absence of CaCl_2_ and in the presence of 2 mM EGTA and 3 mM MgCl_2_. Ratiometric Fura-2AM imaging was performed by in a Nikon Ti-S Inverted Microscope alternatively illuminated with 340 and 380 nm light for 250 ms (Lambda DG-4; Sutter Instrument Co.), using a 20X objective, Nikon CFI Super Fluor. Emission light > 510 nm was captured using a Zyla 4.2 sCMOS camera (Andor). Both channels were collected every 5 seconds, background corrected and analyzed with NIS-Elements software. Images were analyzed in image J.

### Histological analysis

For histological analysis, adipose and liver tissues were fixed in 10% zinc-formalin for 24 h at room temperature and transferred to 70% ethanol for further storage. Tissues were processed, sectioned and stained with hematoxylin and eosin at the Dana Farber Rodent Histopathology facility at Harvard Medical School. Hepatic steatosis scoring was based on the method described in(*40*). Adipose tissue inflammation was evaluated subjectively taking into consideration number and size of inflammatory infiltrates. The inflammatory foci were given a score between 0-3, where 0 represented no inflammation, 1 was moderate, and 3 pronounced was inflammation. Both the liver and adipose tissue were analyzed blindly by 2 independent researchers and by the pathologist from the Dana Farber Rodent Histopathology facility.

### Statistical Analysis

Statistical significance was assessed using GraphPad Prism Version 7. One-way ANOVA (Tukey post-Hoc test) was used for Fig 1D, 2B and 2D. Twoway ANOVA (Sidak post-Hoc test) was used for Fig 4B, 4H and 4I using. Student’s t-tests was used for all other figures. All data is mean ± SEM.

## Acknowledgements

We are grateful to Dr. Andrew Marks from Columbia University, NY for kindly providing us the IP3R1 and IP3R2 floxed mice. We thank Dr. Richard Wojcikiewicz from Upstate Medical University for sending us the IP3R2 antibody generated in his lab. We would like to give special thanks to Dr. Kathryn Claiborn for critical reading and editing of the manuscript. We thank all members of the Sabri Ulker Center and Hotamisligil Lab community for their continued support and encouragement. This work is supported by the Sabri Ulker Center for Metabolic Research. G.P. is supported by an NIH training grant (5T32DK007529-32). E.T. is supported by AHA postdoctoral fellowship (18POST33990109). R.J.D. is supported by NIH R01 DK112698. M.S.H. is supported by AHA Career Development Award (19CDA34660270).

## Author contributions

E.G. and A.P.A. formulated the questions and performed the *in vitro* and *in vivo* experiments, analyzed the data and prepared the figures. G.P., E.C., N.M., E.T., K.I. performed and assisted in vivo experiments. Y.L. and L.G. performed and assisted in vitro experiments. M.S.H. and R.J.D. generated the adipocyte specific JNK1/2 deletion mouse model and performed the JNK1/2 high fat diet studies. G.S.H. and A.P.A conceived and supervised the project, designed experiments, interpreted results, wrote and revised the manuscript.

---

**Supplemental Figure 1:**
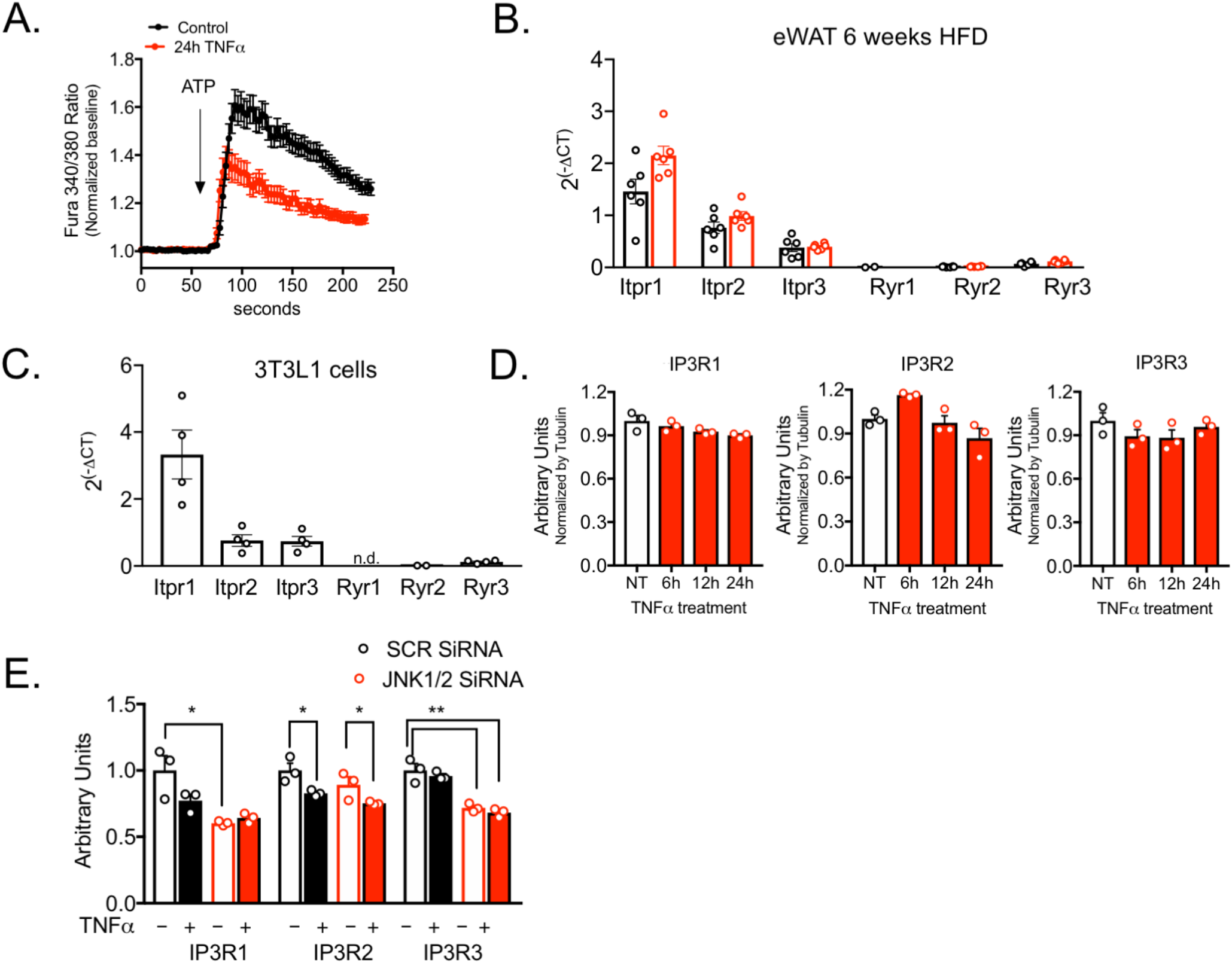
(A) Representative Fura-2AM-based cytosolic Ca^2+^ measurements in 3T3-L1 adipocytes pre-treated with 4ng/mL TNFα or vehicle for 24 hours. ER Ca^2+^ depletion was stimulated with 50 μM of ATP in a media without extracellular Ca^2+^. Representative of 2 experiments. (B) qPCR of mRNA levels of IP3R and Ryanodine R (Ryr) isoforms in adipose tissue derived from lean (LFD) and HFD-fed mice for 6 weeks. Figure show 2^-(ΔCT) comparing gene of interest CT to TBP (control) CT levels. (C) qPCR of mRNA levels of IP3R and Ryanodine R isoforms in 3T3-L1 adipocytes. Figure show 2^-(ΔCT) comparing gene of interest CT to TBP (control) CT levels. (D) Quantification analysis of the western blots presented in Figure 1D, n=3 for each time point. (E) Quantification analysis of the western blots presented in Figure 1F (4 days TNFα treatment).

**Supplemental Figure 2:**
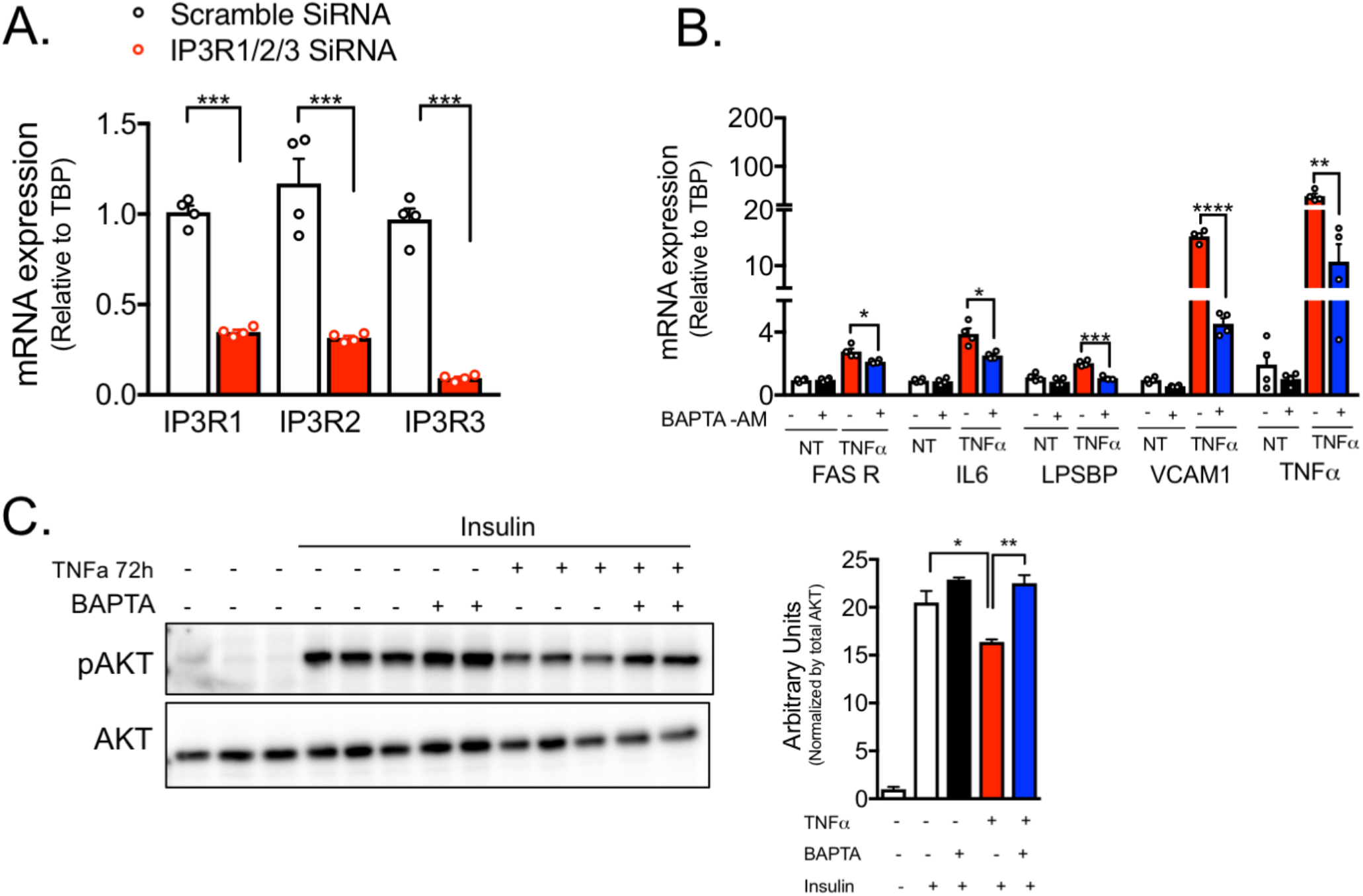
(A) mRNA levels evaluated by qPCR of the indicated genes in 3T3-L1 adipocytes transfected with scrambled (SCR) or IP3R1/2/3 siRNA, n=4 per group, representative of 4 independent experiments, *** p<0.0001. (B) mRNA levels of the indicated genes evaluated by qPCR in 3T3-L1 adipocytes treated with 10 uM of BAPTA-AM in the presence of vehicle or 4ng/mL of TNFα for 6 hours. n=4, representative of 2 independent experiments. *p=0.01 (Fas R, IL6), *** p=0.006 (LPSBP), ****p=0.0004, ** p=0.04. (C) Immunoblot analysis and quantification of insulin signaling after 3 minutes of 3nM insulin treatment in 3T3-L1 adipocytes pre-treated with BAPTA-AM in the presence vehicle or 4ng/mL of TNFα for 72 hours, n=3. In all panels error bars denote s.e.m.

**Supplemental Figure 3:**
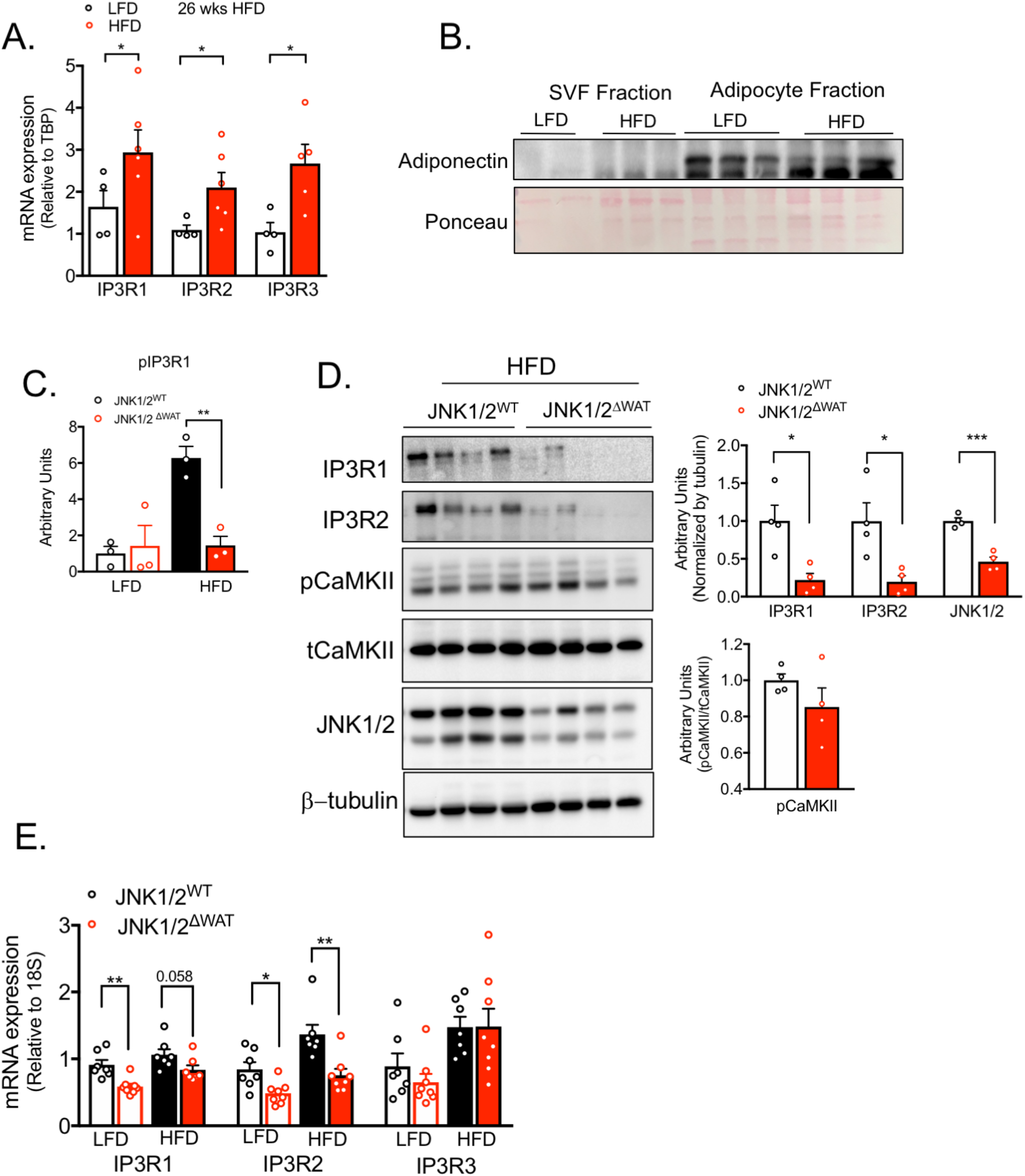
(A) mRNA levels of the indicated genes evaluated by qPCR derived from eWAT of mice fed with low fat diet (LFD) or high fat diet (HFD) for 26 weeks, n=4 for LFD and n=5 for HFD, representative of 2 independent experiments. *p=0.02 (IP3R1, IP3R3), * p=0.04 (IP3R2). (B) Immunoblot analysis of adiponectin expression in stromal vascular fraction (SVF) and adipocyte fraction derived from adipose tissue of animals on LFD and HFD. (C) Quantification of western blot shown in Figure 3F. **p=0.004. (D) Left panel: Immunoblot analysis of protein expression and phosphorylation levels in eWAT derived from controls (JNK1/2^WT^) and adipocyte-specific loss of JNK1/2 (JNK1/2^ΔWAT^) fed LFD or HFD for 16 weeks. Right panel: Quantification of the western blots, n=4 per group, * p<0.01 (IP3R1), *p=0.02, ***p=0.0003 (JNK1/2). (E) mRNA levels of the indicated genes evaluated by qPCR derived from controls (JNK1/2^WT^) and adipocyte-specific loss of JNK1/2 (JNK1/2^ΔWAT^) fed LFD or HFD for 16 weeks. n=7 for JNK1/2^WT^ (LFD and HFD) and n=8 for JNK1/2^ΔWAT^ (LFD and HFD), **p=0.001 (IP3R1), *p=0.01, ** p=0.0029 (IP3R2). In all panels error bars denote s.e.m.

**Supplemental Figure 4:**
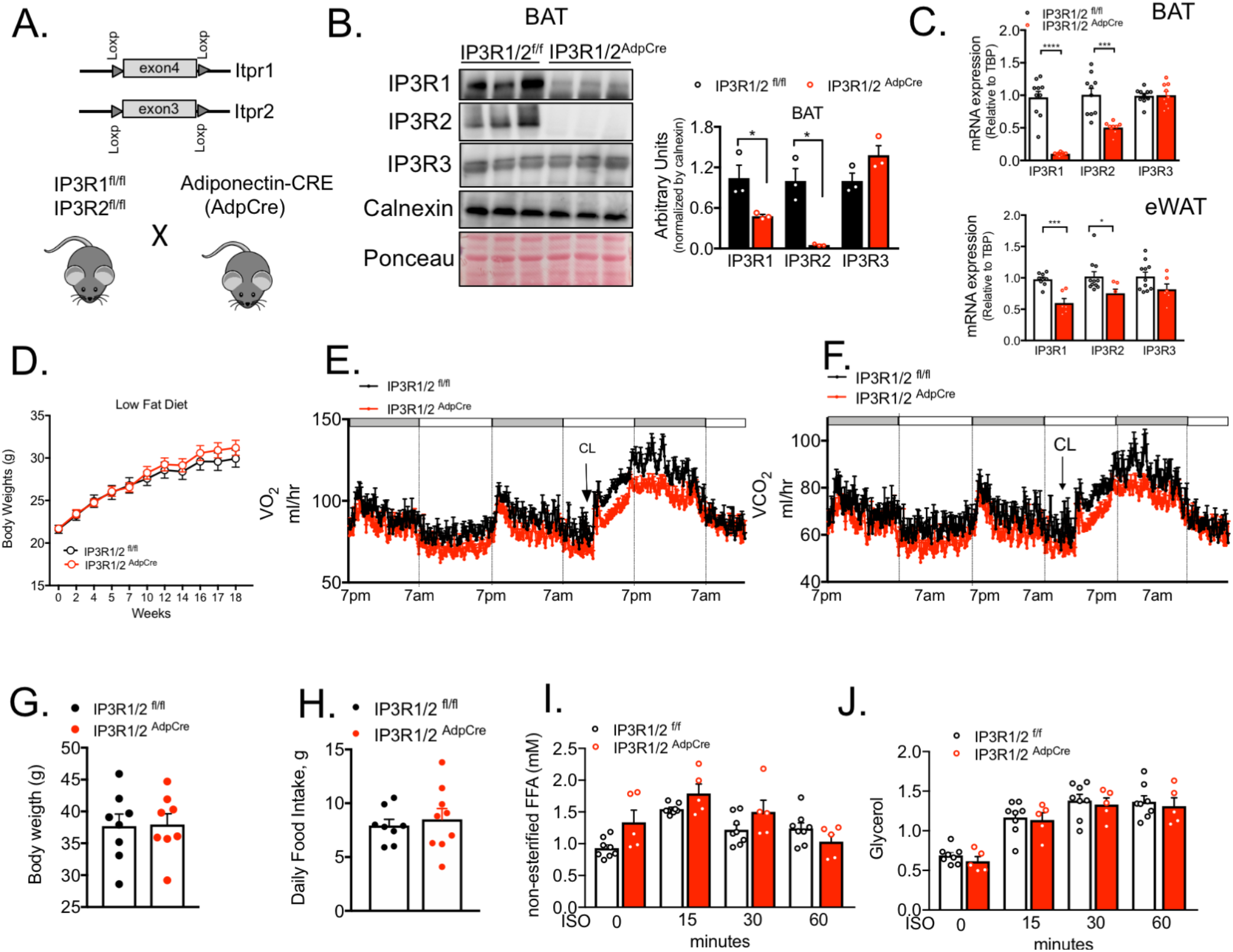
(A) Schematic representation of the animal breeding scheme to obtain mice with adipocyte specific IP3R1/2 deletion. (B) Immunoblot analysis of protein expression levels in brown adipose tissue (BAT) from control (IP3R1/2^fl/fl^ mice) and adipocyte-specific loss of function of IP3R1/2 (IP3R1/2^AdpCRE^) mice on chow diet. Right panel: Quantification of the western blots, n=3 per group, *p<0.042. (C) mRNA levels of the indicated genes evaluated by qPCR from brown adipose tissue (BAT) (upper panel) and white epididymal adipose tissue (eWAT) (lower panel) from control (IP3R1/2^fl/fl^) and adipocyte-specific loss of function for IP3R1/2 (IP3R1/2^AdpCRE^). BAT: n=9 per group, eWAT: n=8 for IP3R1/2^fl/f^ and n=6 for IP3R1/2^AdpCRE^. (D) Weight gain curves of control (IP3R1/2^fl/fl^) and adipocyte-specific loss of function of IP3R1/2 (IP3R1/2^AdpCRE^) mice on low fat diet, n=10 for IP3R1/2^fl/fl^ and n=9 IP3R1/2^AdpCRE^, representative of 3 independent cohorts. (E) Oxygen consumption rates (VO_2_) measured in metabolic cages in control (IP3R1/2^fl/fl^ mice) and adipocyte specific loss of function of IP3R1/2 (IP3R1/2^AdpCRE^) mice on HFD for 8-9 weeks. n= 7 IP3R1/2^fl/f^ and n=8 IP3R1/2^AdpCRE^ per group. (F) Carbon dioxide production rates (VCO2) measured in metabolic cages in control (IP3R1/2^fl/fl^) and adipocyte specific loss of function for IP3R1/2 (IP3R1/2^AdpCRE^) on HFD for 8-9 weeks. n= 7 for IP3R1/2^fl/fl^ and n=8 for IP3R1/2^AdpCRE^. (G) Body weights of the mice presented in E and F. (H) Daily food intake measured during 5 consecutive days in food for control (IP3R1/2^fl/fl^) and adipocyte-specific loss of function of IP3R1/2 (IP3R1/2^AdpCRE^) on HFD for 8-9 weeks. n= 8 per group. (I) Plasma non-esterified free fatty acid (J) Plasma glycerol levels after 6 hours fasting (0 min) and at 15, 30 and 60 minutes after isoproterenol injection (ISO) in control (IP3R1/2^fl/fl^) and adipocyte specific loss of function for IP3R1/2 (IP3R1/2^Ad^p^CRE^) mice fed on HFD for 6 weeks, n= 8 for IP3R1/2^fl/fl^ and n=5 for IP3R1/2^Ad^pc^RE^.

**Supplemental Figure 5:**
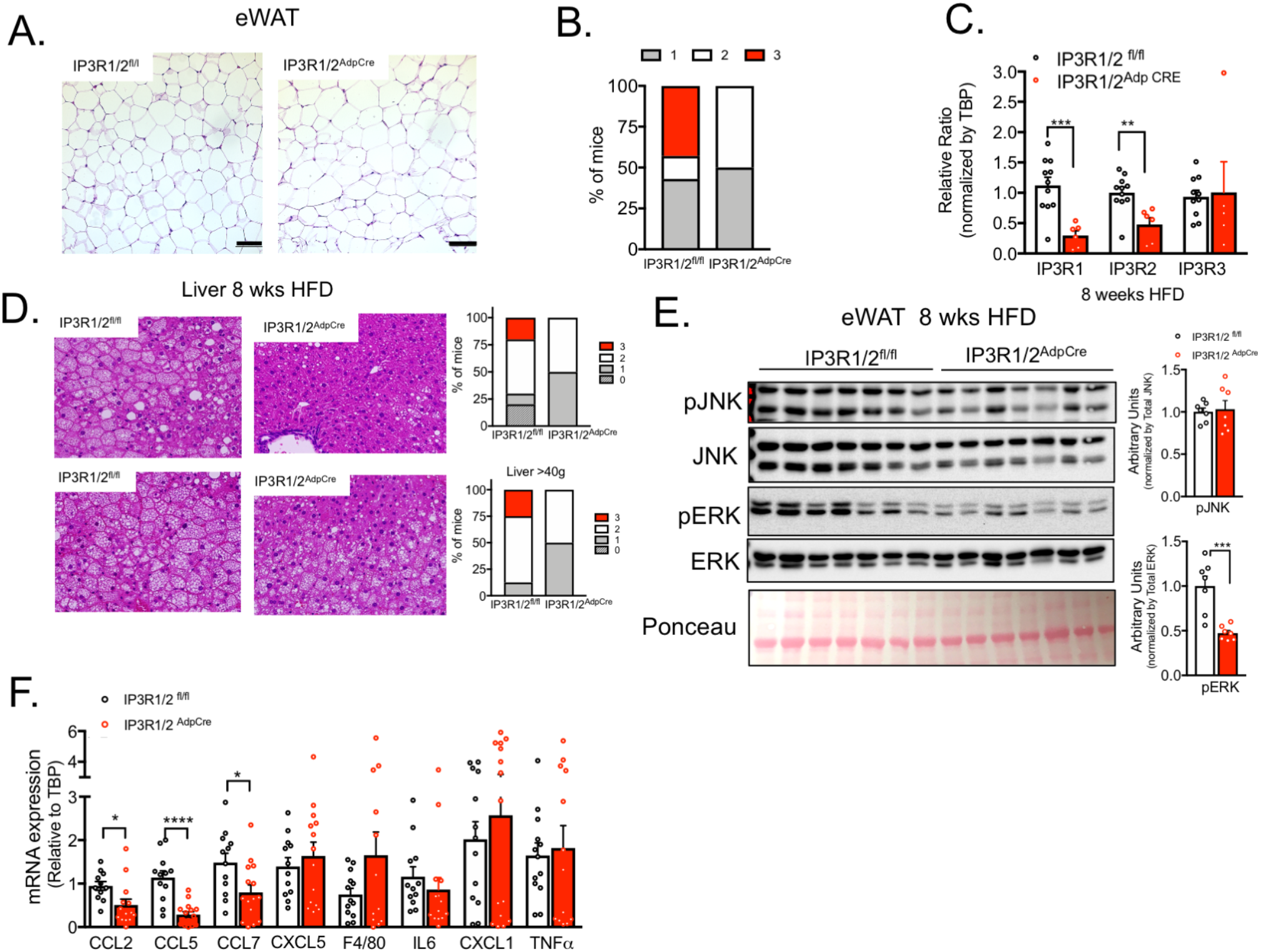
(A) Representative hematoxylin and eosin stained histology section from epidydimal white adipose tissue (eWAT) of control (IP3R1/2^fl/f^) and adipocyte-specific loss of function of IP3R1/2 (IP3R1/2^AdpCRE^) mice fed low fat diet. (B) Scoring of the degree of inflammatory cell infiltration in the tissue. 1-normal or low steatitis, 2-medium steatitis, 3-high steatitis. (C) mRNA levels of the indicated genes evaluated by qPCR derived from eWAT of control (IP3R1/2^fl/f^ mice) and adipocyte-specific loss of function of IP3R1/2 (IP3R1/2^AdpCRE^) mice fed HFD for 8 weeks. (D) Left panel: Representative hematoxylin and eosin stained histology sections from liver of weight-matched control (IP3R1/2^fl/fl^) and adipocyte specific loss of function for IP3R1/2 (IP3R1/2^Ad^p^CRE^) mice fed on HFD for 8-9 weeks, n=11 for IP3R1/2^fl/fl^ and n=6 for IP3R1/2^AdpCRE^. Right panel: Scoring of the degree of hepatosteatosis. 0= normal, 1-low hepatosteatosis, 2-medium hepatosteatosis, 3-high hepatosteatosis. (E) Left panel: Immunoblot analysis of protein expression and phosphorylation levels in epididymal white adipose tissue (eWAT) from control (IP3R1/2^fl/fl^) and adipocyte-specific loss of function of IP3R1/2 (IP3R1/2^AdpCRE^) mice fed on HFD for 8-9 weeks. Right panel: Quantification of the western blots, n=7 per group. *** p<0.001 (F) mRNA levels of the indicated genes evaluated by qPCR derived from eWAT of control (IP3R1/2^fl/fl^) and adipocyte-specific loss of function of IP3R1/2 (IP3R1/2^Ad^p^CRE^) mice fed on HFD for 16 weeks, n= 13 for IP3R1/2^fl/fl^ and 15 for IP3R1/2^Ad^p^CRE^, * p<0.017, ****p<0.0001.

**Supplemental Figure 6:**
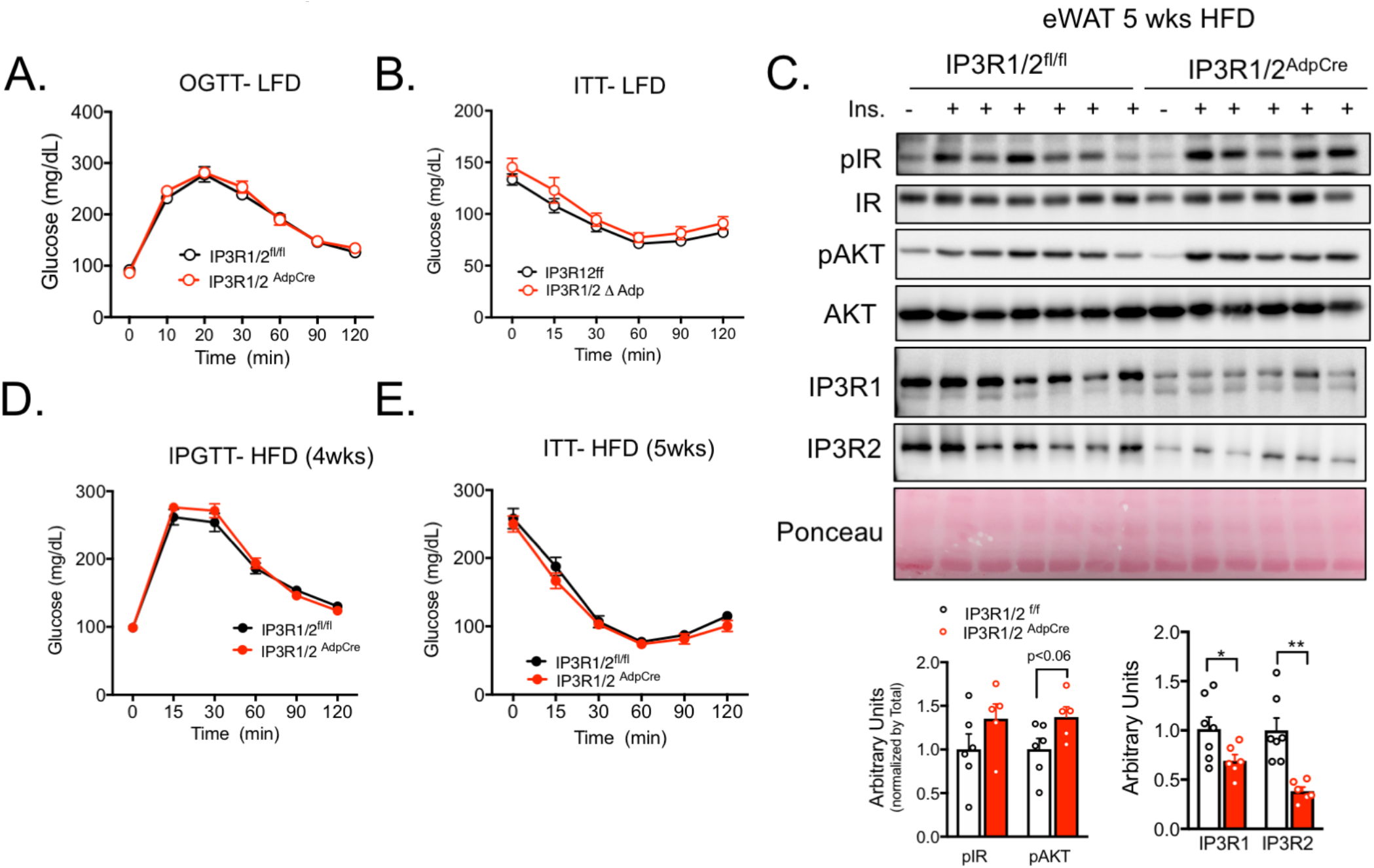
(A) Oral glucose tolerance test in control (IP3R1/2^fl/fl^) and adipocyte-specific loss of function of IP3R1/2 (IP3R1/2^AdpCRE^) fed low fat diet (LFD), n=10 for IP3R1/2^fl/fl^ and n=9 IP3R1/2^AdpCRE^.(B) Insulin tolerance test in control (IP3R1/2^fl/fl^) and adipocyte-specific loss of function of IP3R1/2 (IP3R1/2^AdpCRE^) mice fed on low fat diet (LFD), n=10 for IP3R1/2^fl/fl^ and n=9 for IP3R1/2^Ad^pC^RE^ (C) Upper panel: Markers of *in vivo* insulin signaling evaluated by immunoblot analysis of total eWAT from animals injected with insulin (0.45 U/kg) through the inferior vena cava. Tissues were collected 3 min after injection. Lower panel: phospho-protein level quantification normalized to total protein levels. n= 6 for IP3R1/2^fl/fl^ and n=5 for IP3R1/2^AdpCRE^. (D) Intra-peritoneal glucose tolerance test in control (IP3R1/2^fl/fl^) and adipocyte specific loss of function of IP3R1/2 (IP3R1/2^AdpCRE^) mice fed high fat diet (HFD) for 4 weeks, n=9 for IP3R1/2^fl/fl^ and n=10 for IP3R1/2^AdpCRE^. (E) Insulin tolerance test in control (IP3R1/2^fl/fl^) and adipocyte-specific loss of function of IP3R1/2 (IP3R1/2^Ad^p^CRE^) mice fed HFD for 5 weeks, n=9 for IP3R1/2^fl/fl^ and n=10 for IP3R1/2^Ad^p^CRE^.

**Supplemental Table 1.**
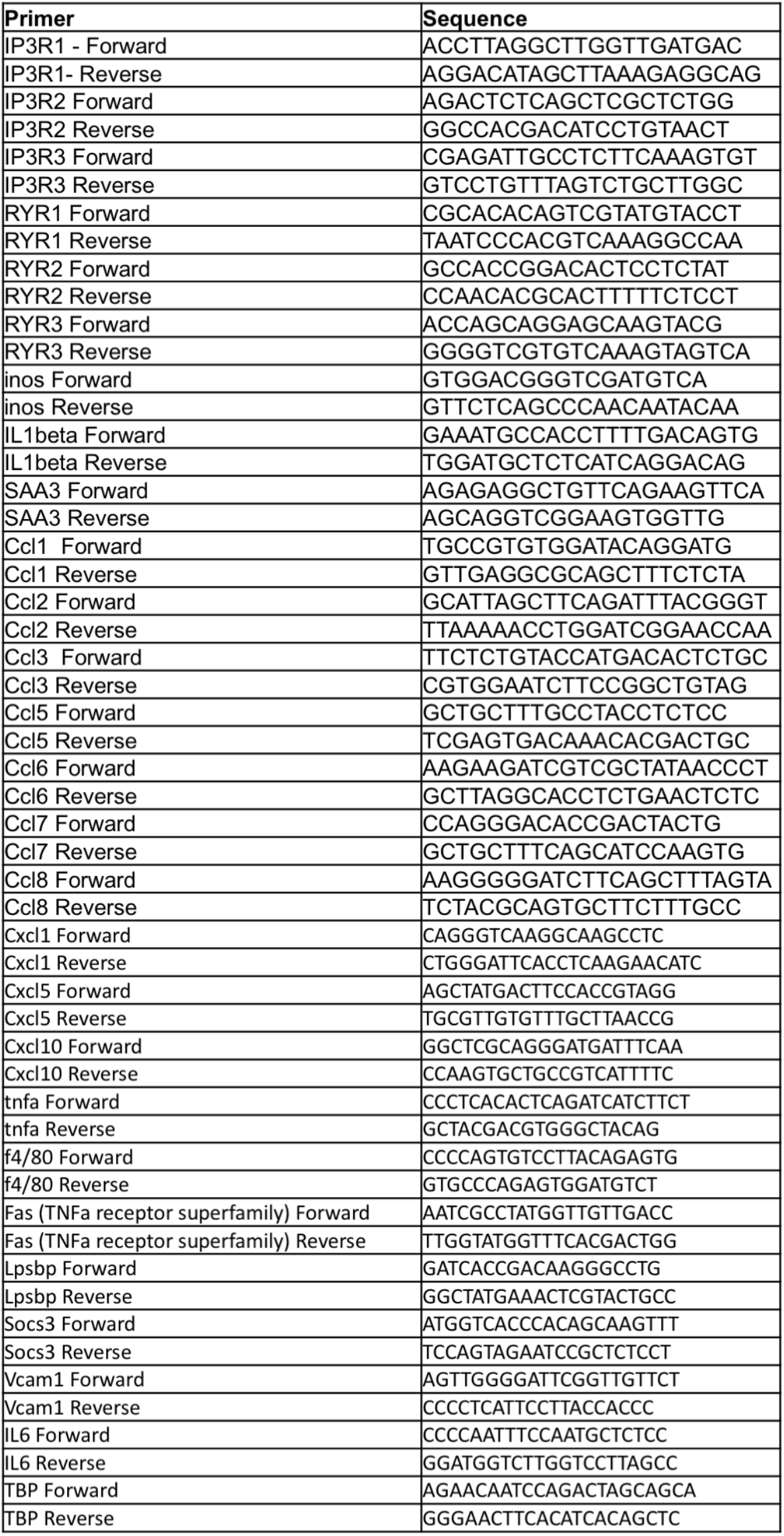

